# Gene expression tradeoffs determine bacterial survival and adaptation to antibiotic stress

**DOI:** 10.1101/2024.01.20.576495

**Authors:** Josiah C. Kratz, Shiladitya Banerjee

**Affiliations:** Computational Biology Department, Carnegie Mellon University, Pittsburgh, PA 15213, USA; Department of Biological Sciences, Carnegie Mellon University, Pittsburgh, PA 15213, USA; Department of Physics, Carnegie Mellon University, Pittsburgh, PA 15213, USA

## Abstract

To optimize their fitness, cells face the crucial task of efficiently responding to various stresses. This necessitates striking a balance between conserving resources for survival and allocating resources for growth and division. The fundamental principles governing these tradeoffs is an outstanding challenge in the physics of living systems. In this study, we introduce a coarse-grained theoretical framework for bacterial physiology that establishes a connection between the physiological state of cells and their survival outcomes in dynamic environments, particularly in the context of antibiotic exposure. Predicting bacterial survival responses to varying antibiotic doses proves challenging due to the profound influence of the physiological state on critical parameters, such as the Minimum Inhibitory Concentration (MIC) and killing rates, even within an isogenic cell population. Our proposed theoretical model bridges the gap by linking extracellular antibiotic concentration and nutrient quality to intracellular damage accumulation and gene expression. This framework allows us to predict and explain the control of cellular growth rate, death rate, MIC and survival fraction in a wide range of time-varying environments. Surprisingly, our model reveals that cell death is rarely due to antibiotic levels being above the maximum physiological limit, but instead survival is limited by the inability to alter gene expression sufficiently quickly to transition to a less susceptible physiological state. Moreover, bacteria tend to overexpress stress response genes at the expense of reduced growth, conferring greater protection against further antibiotic exposure. This strategy is in contrast to those employed in different nutrient environments, in which bacteria allocate resources to maximize growth rate. This highlights an important tradeoff between the cellular capacity for growth and the ability to survive antibiotic exposure.

## I. INTRODUCTION

Bacteria must regularly cope with a diverse set of harsh environments in their natural habitats. In unpredictable conditions, cells must balance the competing objectives of replication and protection against stress. Previous work has focused on identifying the molecular players which control the bacterial stress response and understanding how specific genes confer protection against specific stressors [1–4]. In addition, the role of phenotypic heterogeneity and population-level bet hedging strategies in the bacterial stress response have been studied [2, 5–7]. However, the fundamental principles governing the tradeoffs between expression of these genes and genes needed for growth is an outstanding question.

Antibiotic exposure is one such pertinent environmental stressor. Antibiotics are often produced by competing microbes [8, 9], and are commonly used in the treatment of human infections [10]. Systems-level changes to bacterial physiology induced by antibiotic exposure, such as changes to cellular growth rate [11–14], gene expression [6, 15–17], and cell morphology [18–21] have been well-characterized. As a result, much is known about the proximate causes of antibiotic action, but vastly less is known about how these causes ultimately lead to bacterial cell death, and how cell death is abated by systems-level changes to cell physiology [22]. Furthermore, killing efficiency is not solely dependent on antibiotic dose, but on many other factors including the environment and the physiological state of the cell (Figure 1A). As such, to understand bacterial stress response strategies and to predict antibiotic efficacy in different environments, it is necessary to link environment not only to growth physiology, but also to damage accumulation and cell viability.

**FIG. 1.**
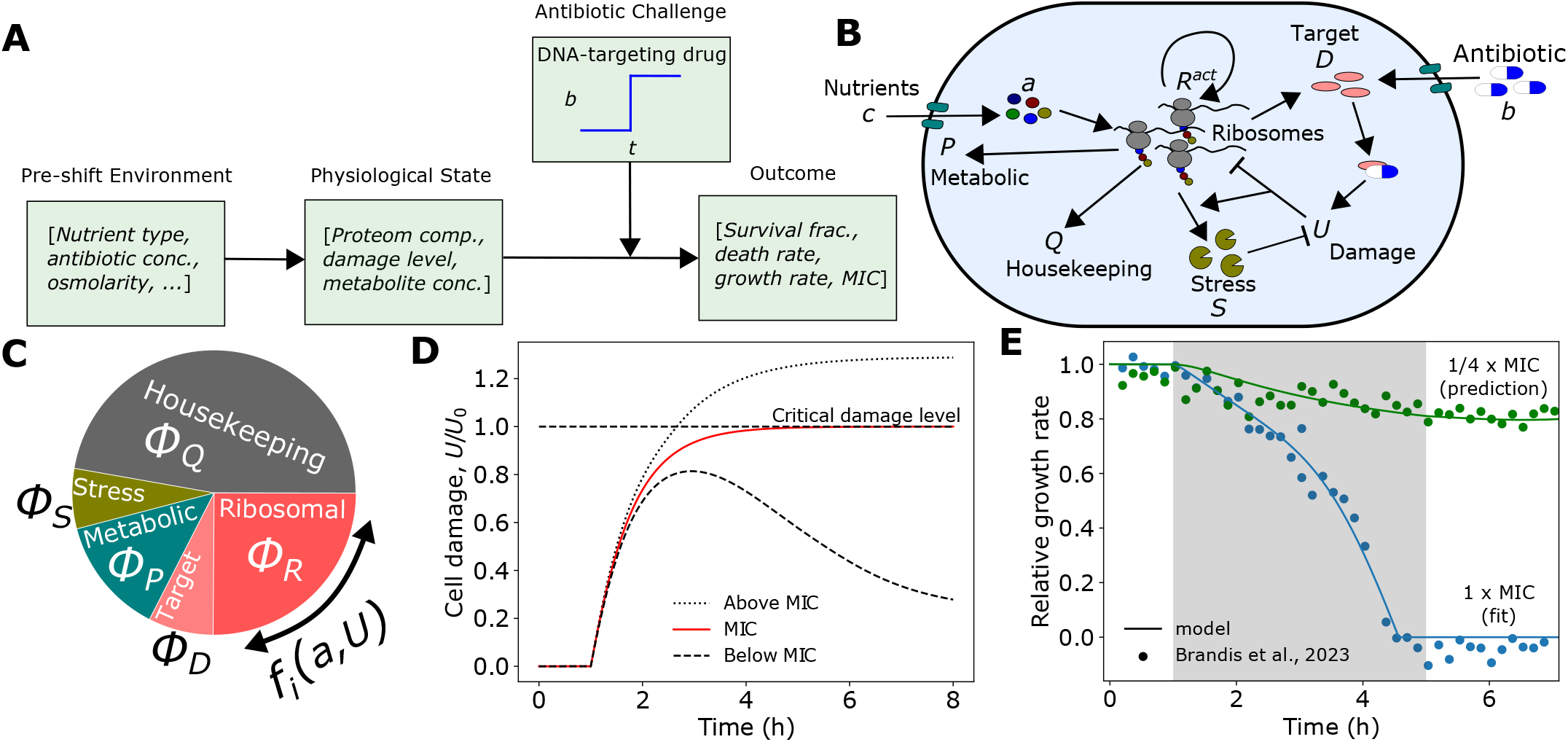
Coarse-grained model for cell growth and death in dynamic antibiotic environments. (A) The survival outcome of a bacterial population exposed to antibiotics is heavily dependent on the pre-shift environment through its influence on the physiological state of the cell. (B) Schematic of coarse-grained model of bacterial physiology. Nutrients (*c*) are imported by metabolic proteins (P) and converted to amino acids (*a*), which are then consumed by ribosomes (R) to produce proteins. Antibiotics (*b*) enter the cell and bind their intracellular target (D) to produce damage which can be repaired by stress proteins (S). (C) By dynamically regulating the fraction of the total translational flux devoted to each proteome sector *i, fi*, in response to changes in *a* and *U* triggered by environmental changes, the cell alters its proteome composition, thus altering its susceptibility to further antibiotic challenge. (D) In our model framework, the Minimum Inhibitory Concentration (MIC) is defined as the minimum value of *b* which causes *U* to cross the critical damage threshold (*U* = *U*_0_). (E) Model successfully explains antibiotic-induced growth reductions for different values of *b*. Grey region indicates antibiotic application. Experimental data are of *E. coli* BW25993 cells in LB exposed to 8 (green) and 32 (blue) *µ*g/ml of Ciprofloxacin from Ref. [14] (see Fig. S3 for data analysis details). See Table 1 for a list of model parameters, and Fig. S5 for the effects of parameter variations on growth rate dynamics

Previous work has shown that death rate increases approximately linearly with growth rate, but that the sensitivity of death rate to changes in growth rate depends significantly on environment and metabolic state [17, 23–25] Many mathematical models have been developed to link antibiotic dose to growth rate [13, 26–28] but little has been done to connect growth physiology mechanistically to cell survival outcome. A recent work [17] identified a general stress-response sector in *E. coli*, whose expression reduces death rate. However, many questions remain unanswered: how is resource allocation to stress protein production mechanistically linked to the environment, and what specific effects does it have on cell physiology to mitigate antibiotic-induced death?

**TABLE I.** Model parameters. See Appendix A for more details.

| Parameter | Description | Value | Figure number |
| --- | --- | --- | --- |
| $\phi_R^{\max}$ | maximum flux allocation to ribosome production [12] | 0.55 | all |
| $a_t$ | translation attenuation threshold [45] | $10^{-4}$ | all |
| $a_n$ | feedback inhibition threshold [45] | $10^{-3}$ | all |
| $\kappa_t^0$ ($\text{h}^{-1}$) | translational efficiency rate constant, strain specific, fitted | 2.7 | 2-6 |
|  |  | 3.5 | 1 |
| $\kappa_n^0$ ($\text{h}^{-1}$) | nutritional efficiency rate constant, nutrient specific, calculated | 5.5 | 2-6 |
|  |  | 1.12 | 1 |
| $\alpha/U_0$ ($\frac{\text{ml}}{\mu\text{g h}}$) | normalized damage production rate constant, nalidixic acid, fitted | 1.54 | 2-4, 5D, 6 |
|  | normalized damage production rate constant, ciprofloxacin, fitted | 0.13 | 1, 5A-C |
| $\beta$ ($\text{h}^{-1}$) | damage removal rate constant, nalidixic acid, fitted | 10.5 | 2-4, 5D, 6 |
|  | damage removal rate constant, ciprofloxacin, fitted | 5.29 | 1, 5A-C |
| $K_u$ | value of half-maximal expression, strain specific, fitted | 0.076 | 2-6 |
|  |  | 0.030 | 1 |
| $\phi_S^{\max}$ | maximum flux allocation to stress protein production, antibiotic specific, fitted | 0.33 | 2-4, 5D, 6 |
|  |  | 0.18 | 1, 5A-C |
| $\sigma$ | noise amplitude, fitted | 0.015 | 5, 6 |
| $\mu_X$ ($\text{h}^{-1}$) | division protein degradation rate [40] | 0.6 | 5, 6 |
| $\gamma_\alpha$ ($\mu\text{m}^{-3}$) | division protein production parameter [40] | 4.5 | 5, 6 |
| $\gamma_\beta$ ($\mu\text{m}^{-3}$) | division protein production parameter [40] | 1.1 | 5, 6 |

**TABLE II.**
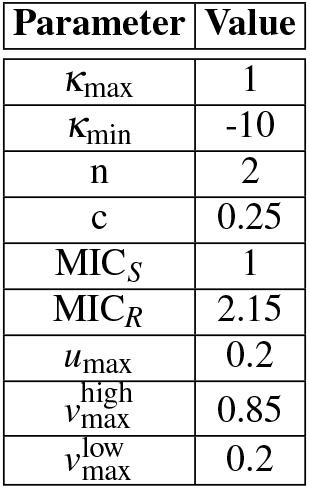
Model parameters for phenotypic switching model.

| Parameter | Value |
| --- | --- |
| $\kappa_{\max}$ | 1 |
| $\kappa_{\min}$ | -10 |
| n | 2 |
| c | 0.25 |
| $\text{MIC}_S$ | 1 |
| $\text{MIC}_R$ | 2.15 |
| $u_{\max}$ | 0.2 |
| $v_{\max}^{\text{high}}$ | 0.85 |
| $v_{\max}^{\text{low}}$ | 0.2 |

To gain a systems-level understanding of how cellular stress response and growth are connected to the environment and antibiotic killing efficiency, we have developed a multi-scale model for cell growth and death which coarse-grains cellular physiology into a limited number of state variables and kinetic parameters to predict both single-cell and population-level behavior. Specifically, our model connects extracellular antibiotic concentration and nutrient quality to the stochastic dynamics of damage accumulation and proteome allocation to predict bacterial growth rate, death rate, and survival fraction in a wide range of time-varying environments.

We apply our model to predict changes in Minimum Inhibitory Concentration (MIC) of antibiotics as a function of the environment in response to replication-targeting bactericidal antibiotics. We find that cells with reduced growth rates caused by stressful pre-shift environments are able to survive higher concentrations of antibiotics (increased MIC), in agreement with recent experimental data [17]. Our model predicts. that this non-intuitive relationship between growth and death is a consequence of the dynamics of damage accumulation and removal, which are heavily dependent on the initial physiological state of the cell. Specifically, cells which are pre-exposed to low levels of antibiotics overexpress genes that can repair antibiotic-induced damage. Thus, when exposed to higher levels of antibiotics, they can more quickly repair new damage and survive, despite starting with an initially higher level of damage.

Our model predicts that there is a maximum antibiotic dose above which bacteria cannot survive unless through mutation, regardless of physiological state. However, model analysis reveals that cell death is rarely due to antibiotic levels reaching this limit, but instead survival is limited by the inability of a cell to alter gene expression sufficiently quickly to transition to a less susceptible physiological state. Our model highlights a critical gene expression tradeoff between growth and survival: allocation to stress response pathways is imperative to survive antibiotic challenge, but investment in these pathways reduces the resources available for growth. Thus, our model predicts that non-growth optimal proteome allocation increases bacterial survival compared to growth optimal allocation, a strategy which is preferred in many nutrient environments [29–31]. This tradeoff between allocation towards growth and stress response provides an explanation for the non-growth optimal allocation observed in *E. coli* [11, 15].

## II. RESULTS

### A. Resource allocation theory of cellular stress response in dynamic environments

Stress-induced cell death can be a consequence of many factors. For bactericides in particular, cell death is not simply a result of target-specific inhibition. Instead, primary drug-target interactions perturb various metabolic pathways to induce an array of downstream effects, which can cause damage to both DNA and proteins [22, 32–3 To combat such damage, bacteria can induce both a nonspecific and specific stress response in which many similar proteins are up-regulated in response to nutrient, antibiotic, or osmotic stress [17, 34, 36, 37]. Bactericide-induced damage can be repaired by such proteins, e.g. SOS proteins in the case of DNA damage [1], allowing bacteria to survive and grow despite antibiotic challenge. Importantly, over short timescales survival is mediated by changes in gene expression, and is not due to genetic mutations [17, 38].

Here we model physiological effects of bactericidal antibiotics, specifically those targeting DNA replication. Motivated by the common mechanism of cellular death induced by bactericides, we propose a coarse-grained model of damage accumulation and removal and connect it to cell physiology to predict bacterial growth and survival (Figure 1B). Specifically, the dynamics of the damage concentration *U* carried by a single cell can be expressed as

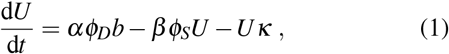

where *b* is the antibiotic concentration, which produces damage at a concentration specific rate *α* when bound to its target protein *D*, e.g. DNA gyrase in the case of quinolones. Here *ϕ*_*D*_ represents the mass fraction of *D*, and *U* represents the total concentration of damage incurred by a single cell, which may include factors like misfolded proteins, membrane and DNA damage, or other contributors to cell death. This coarse-grained approach to modeling cell damage has recently proved successful in the context of bacterial aging [39]. Damage is actively removed by stress proteins *S*, with mass fraction *ϕ*_*S*_, at a rate *β*, and is also diluted with growth rate *κ*. Cell death occurs when damage accumulation exceeds a critical level *U*_0_, such that *U* (*t* = *τ*_death_) = *U*_0_. Mathematically, this threshold is the value of *U* above which the fixed point of the dynamical system, corresponding to survival, is no longer accessible in the deterministic model (discussed in more detail is section II.C). We assume that the dynamics of damage accumulation are much slower than the dynamics of antibiotic import and target binding, and thus model changes in *b* as instantaneous. Critically, bacteria alter *ϕ*_*S*_ and *ϕ*_*D*_ in response to environ-mental changes. Thus, we connect damage accumulation dynamics to changes in gene expression following our recently introduced framework for dynamic proteome allocation [40]. In brief, cells import and convert nutrients to amino acids, with mass fraction *a*, via metabolic proteins, with protein mass fraction *ϕ*_*P*_. Amino acids are consumed by translating ribosomes, with mass fraction *ϕ*_*R*_, to synthesize all proteins, including themselves. As a result, *ϕ*_*R*_ sets the cellular growth rate, specifically *κ* = *κ*_*t*_(*a,U*)*ϕ*_*R*_, where *κ*_*t*_(*a,U*) is the translational efficiency. Importantly, the translational efficiency is reduced under conditions of limited amino acid availability and elevated damage levels, in order to capture the effects of damage on the translational machinery (see Appendix A for details). The dynamics of each sector are given by

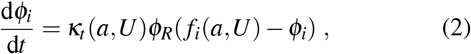

where *i* = [*P, R, S, D, Q*] and *f*_*i*_(*a,U*) denotes the fraction of total cellular protein synthesis flux devoted to sector *i*, and can be a function of *a* and/or *U*. We impose two constraints on the model motivated by *E. coli* proteomics data. First, a significant portion of the proteome is invariant to environmental perturbations [41], thus we define the housekeeping sector such that *ϕ*_*Q*_ = *f*_*Q*_ = constant, and

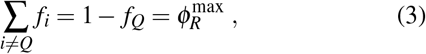

*R* where 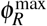 is the upper limit to the allocation fraction devoted to ribosomal proteins. Second, steady-state proteomics data [16] revealed that the molecular targets of many antibiotics which inhibit DNA replication, such as DNA gyrases, are coregulated with ribosomal proteins under carbon, nitrogen, and translation limiting regimes (Fig S1). As this work focuses on replication-targeting bactericides, we assume that the target sector, *ϕ*_*D*_, is coregulated with the R sector, such that *ϕ*_*D*_ ∝ *ϕ*_*R*_. These constraints reduce the number of independent sectors to two, namely *ϕ*_*R*_ and *ϕ*_*S*_.

Cells exhibit a general stress response which is induced in response to cellular damage, and is mediated by various signaling molecules and transcription factors including ppGpp and RpoS [1, 36, 37, 42, 43]. Thus, *f*_*S*_ is indirectly activated by *U*, and we model its dependence by a simple sigmoidal function, 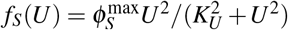, where expression sat-urates at 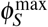 and *K*_*U*_ is a constant. Additionally, steady-state transcriptomic analysis revealed that ribosomal sector expression is reduced to allow for stress sector expression [17]. As such, the fraction of total synthesis capacity devoted to ribosomes, *f*_*R*_, is now a function of both *a* and *U*, where the maximum value of *f*_*R*_ is reduced as *U* increases (see Appendix A for details). When *U* = 0, *f*_*R*_ is chosen to maximize translational flux at steady-state, thus maximizing growth rate [40, 44, 45].

Lastly, the dynamics of the amino acid mass fraction are given by the difference in the metabolic and translational fluxes, specifically

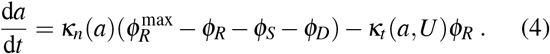

Our model now has two key kinetic variables: *a* and *U* (Figure 1C). *a* acts as a readout of flux imbalance, driving metabolic and ribosomal proteome reallocation in response to nutrient changes. Increase in *U* caused by antibiotic application drives stress protein expression, which in turn can impact allocation to the other sectors.

*Predicting Minimum Inhibitory Concentration*. – Our model can be utilized to predict the Minimum Inhibitory Concentration (MIC), typically defined as the antibiotic concentration threshold beyond which a bacterial population can experience complete extinction, while concentrations below the MIC allow the population to persist [38, 46]. Thus, for a given pre-shift environment, the MIC can be predicted using our framework by identifying the minimum post-shift antibiotic concentration *b* which results in cell damage accumulating to the critical threshold, *U* (*t*) = *U*_0_ (Figure 1D), which corresponds to cell death. This value can be obtained by solving the constrained optimization problem:

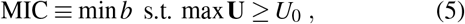

where **U** denotes the vector of damage values across time (see Appendix B for more details).

Taken together, Eqs. 1-4 define our model. This model can be fit well to experimental data, and yields extremely accurate predictions for growth rate dynamics for other antibiotic concentrations above and below the MIC not used in fitting (Figure 1E and Fig S2A). Critically, when antibiotics are removed, growth rate recovers to its pre-shift value for antibiotic concentrations below the MIC, but does not recover for concentrations above the MIC (Fig S2B).

### B. Growth rate control under stress

Bacteria must quickly alter gene expression to adapt to environmental stress. Our model can be utilized to predict the dynamics of damage accumulation, proteome allocation, and growth rate in response to time-varying antibiotic stress (Figure 2A). Antibiotic application leads to accumulation of damage, causing a sharp increase in allocation to stress protein production. Allocation to stress proteins largely comes at the expense of ribosomal allocation. This reduction in *ϕ*_*R*_, in combination with the increase in *U*, results in a growth rate reduction (Figure 2A).

**FIG. 2.**
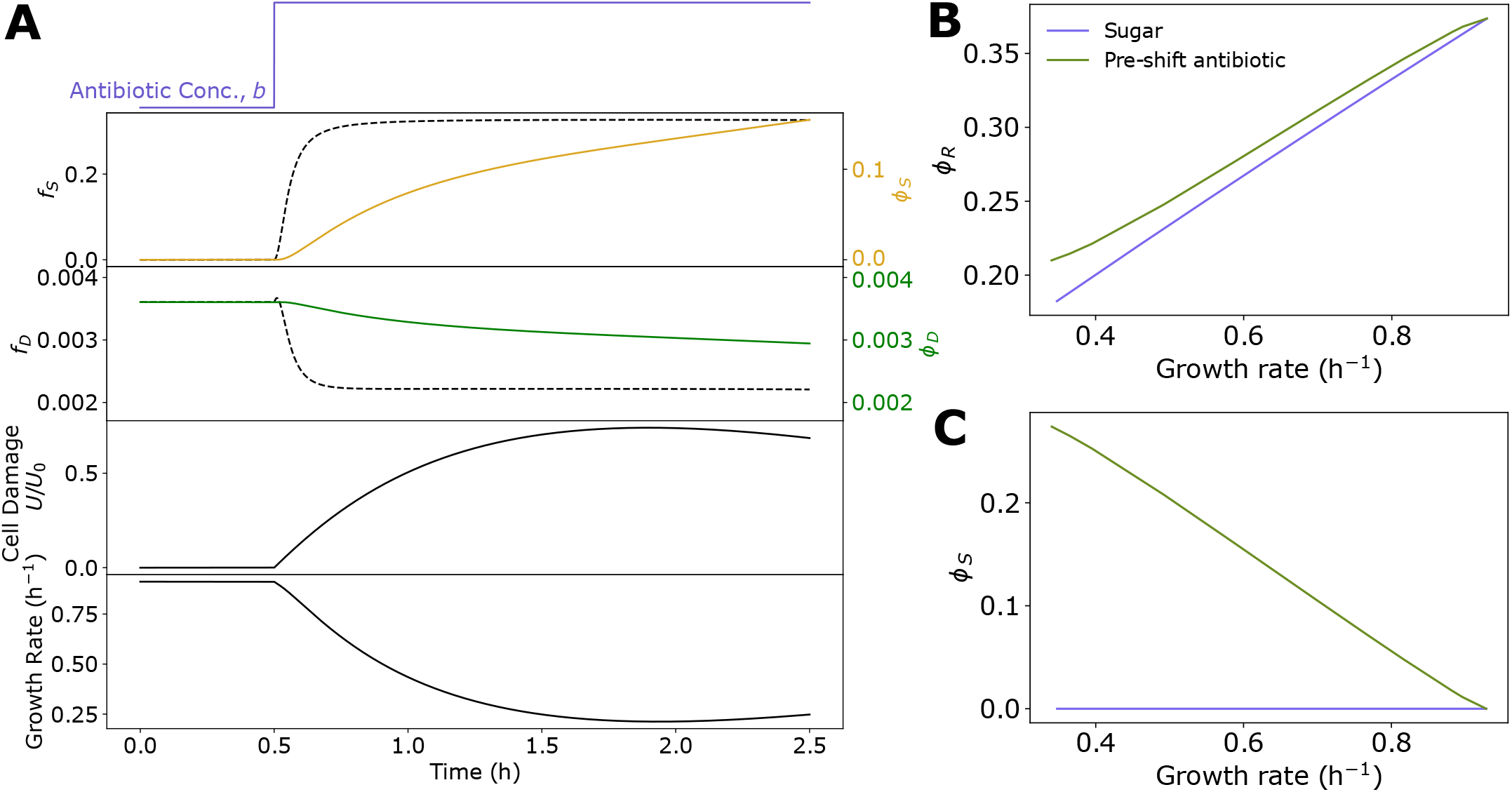
Proteome reallocation and damage accumulation predicts bacterial growth rate across stress conditions. (A) Top to bottom: antibiotic concentration, stress sector mass fraction, antibiotic target sector mass fraction, cell damage, and growth rate (*κ*) dynamics in response to stepwise application of replication-targeting bactericide at *t* = 1h. (B,C) Ribosome (B) and stress (C) sector mass fraction as a function of growth rate for different nutrient environments (blue) and for different concentrations of pre-shift antibiotic exposure (green). See Table 1 for a list of parameters.

Furthermore, the model is able to qualitatively capture experimentally-observed [17] relationships between proteome allocation and growth rate across both nutrient and antibiotic conditions. Specifically, the model predicts that ribosomal sector allocation decreases with growth rate when growth is reduced either through a reduction in nutrient quality or through an increase in antibiotic concentration (Figure 2B). In contrast, stress sector expression is not affected by changes to nutrient quality which reduce growth rate, but sector allocation increases with decreasing growth rate when growth is reduced by increasing the applied antibiotic concentration (Figure 2C).

### C. Effect of pre-shift environment on MIC

Interestingly, when comparing the relationship between pre-shift growth rate and MIC, the model predicts very different behavior based on the pre-shift environment (Figure 3A). Decreasing the growth rate by decreasing the nutrient quality has little impact on the MIC. However, decreasing the growth rate by exposure to low levels of pre-shift antibiotic result in significant increases in MIC, with higher pre-shift doses resulting in higher MIC values. These predictions quantitatively capture recent experimental results [17], and highlight the important role the environment plays in determining bacterial fitness in response to antibiotic challenge.

**FIG. 3.**
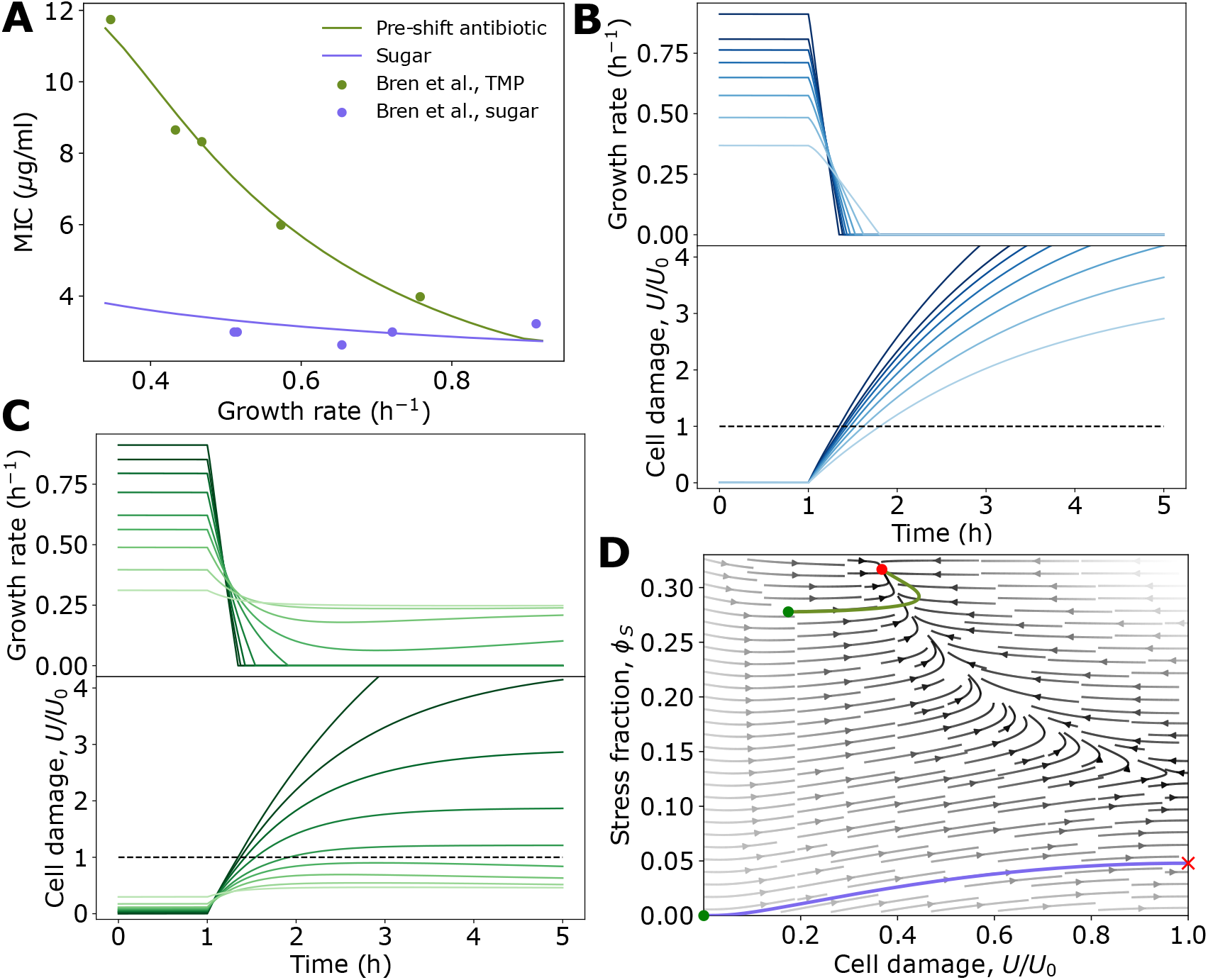
Pre-shift environment heavily influences damage accumulation dynamics and survival. (A) Minimum Inhibitory Concentration (MIC) as a function of pre-shift growth rate under nutrient limitation (blue), and growth inhibition via low levels of pre-shift antibiotic (green), as computed from (5). Experimental data are from from Ref. [17] and are of *E. coli* NCM3722 cells grown in different sugars or exposed to different amounts of the bactericidal antibiotic TMP, before the MIC of nalidixic acid was measured. Model behavior is robust to parameter choice (see Fig. S6). (B,C) Example trajectories of growth rate and damage accumulation for decreasing nutrient quality (B) and increasing pre-shift antibiotic concentration (C). (D) Phase portrait in *ϕ*_*S*_-*U* plane with two example trajectories corresponding to pre-shift antibiotic exposure (green) and no exposure (blue). Green dots denote starting position, and red dot and X denotes fixed point and cell death, respectively. See Table 1 for a list of parameters.

Our model predicts that these fitness gains by bacteria pre-exposed to antibiotics are explained by the differences in proteome allocation and their impact on the dynamics of damage accumulation. Cells in different nutrient environments initially do not carry damage, and so the stress sector is not expressed (Figure 2C and Figure 3B). As a result, when antibiotics are applied, cells quickly accumulate damage regardless of initial growth rate, resulting in very similar values of MIC (Figure 3A,B). This can be seen in (1), where when *ϕ*_*S*_ is small, the dynamics of *U* are largely dictated by *b* (the smaller impact of *ϕ*_*D*_ will be discussed in later sections). In contrast, cells exposed to increasing initial levels of antibiotic both have increasing initial damage levels (Figure 3C), but also have increased stress sector expression (Figure 2C). This yields the counterintuitive result that cells which initially have more damage are able to withstand higher antibiotic concentrations, resulting in an increased MIC (Figure 3A,C). Again this can be explained in terms of the dynamics of *U* : Slower growing cells have higher initial values of *ϕ*_*S*_, thus when additional antibiotics are applied, the damage removal rate is significantly higher, resulting in a higher value of *b* required for *U* to go above *U*_0_.

An important prediction of this model is that the dynamics of damage accumulation and removal dictate survival, and that these dynamics are heavily influenced by the initial physiological state of a cell. Consequently, cell fate can be determined by considering a cell’s position in the *ϕ*_*S*_-*U* phase space immediately before an antibiotic shift. For most environments, there exists only one positive real steady-state solution of the system for a given antibiotic concentration, *b*, corresponding to a stable fixed point which describes the steady-state physiology of surviving bacteria (growth bistability is predicted to occur only in very poor nutrient and high antibiotic environments, see Appendix C). Critically, this fixed point is not accessible from all regions of phase space. The trajectories of cells characterized by high damage levels and/or low stress sector expression will diverge away from the fixed point, resulting in cell death when *U* = *U*_0_ (Figure 3D). Only cells with elevated values of *ϕ*_*S*_ and/or low values of *U* will have trajectories which arrive at the fixed point, corresponding to survival (Figure 3D).

Above a threshold antibiotic concentration *b*_max_, all fixed points become imaginary. This transition corresponds to the maximum survivable antibiotic concenation without mutation, and is given by 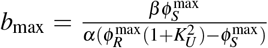 This value is much higher than concentrations typically required to cause cell death, as this fixed point is almost entirely inaccessible. Thus, our model reveals that cell death is rarely due to antibiotic levels being above the maximum physiological limit, but instead survival is limited by the inability to transition to the appropriate physiological state.

### D. Non-growth optimal resource allocation promotes bacterial survival

As shown in the last section, a cell’s survival is heavily dependent on the pre-shift environment, i.e., its initial position in the *ϕ*_*S*_-*U* phase space. As a result, the specific way bacteria allocate resources in response to antibiotic exposure, and thus alter their position in *ϕ*_*S*_-*U* space, has a significant impact on surviving additional increases in antibiotic level. For this reason, we were interested in comparing our proposed model to other potential resource allocation strategies. Bacteria are known to allocate resources to maximize growth rate in many different nutrient environments [29, 44, 45], and so we compared our model to the growth optimal strategy.

The growth optimal strategy was implemented by computing the growth optimal proteome allocation (subject to the constraints of (3)) for each environment, using these values to set the allocation fractions *f*_*i*=[*P,R,S,D*]_. Figure 4A shows stress protein expression as a function of damage level for both resource allocation strategies. Evidently, our proposed model significantly over-expresses stress proteins compared to the growth optimal model for low and moderate damage levels. This result is consistent with experimental results showing that *E. coli* resource allocation is not growth-rate optimal when exposed to replication-targeting bactericides [11], but also raises the question, are there any potential fitness advantages conferred by this non-optimal resource allocation strategy? To answer this, we computed the predicted MIC values for the growth optimal strategy in a range of pre-shift antibiotic environments. In all environments tested, the MIC is significantly reduced compared to the non-growth optimal model (Figure 4B). Thus, our model suggests that cells trade reduced growth for increased survival chances in bactericidal environments.

**FIG. 4.**
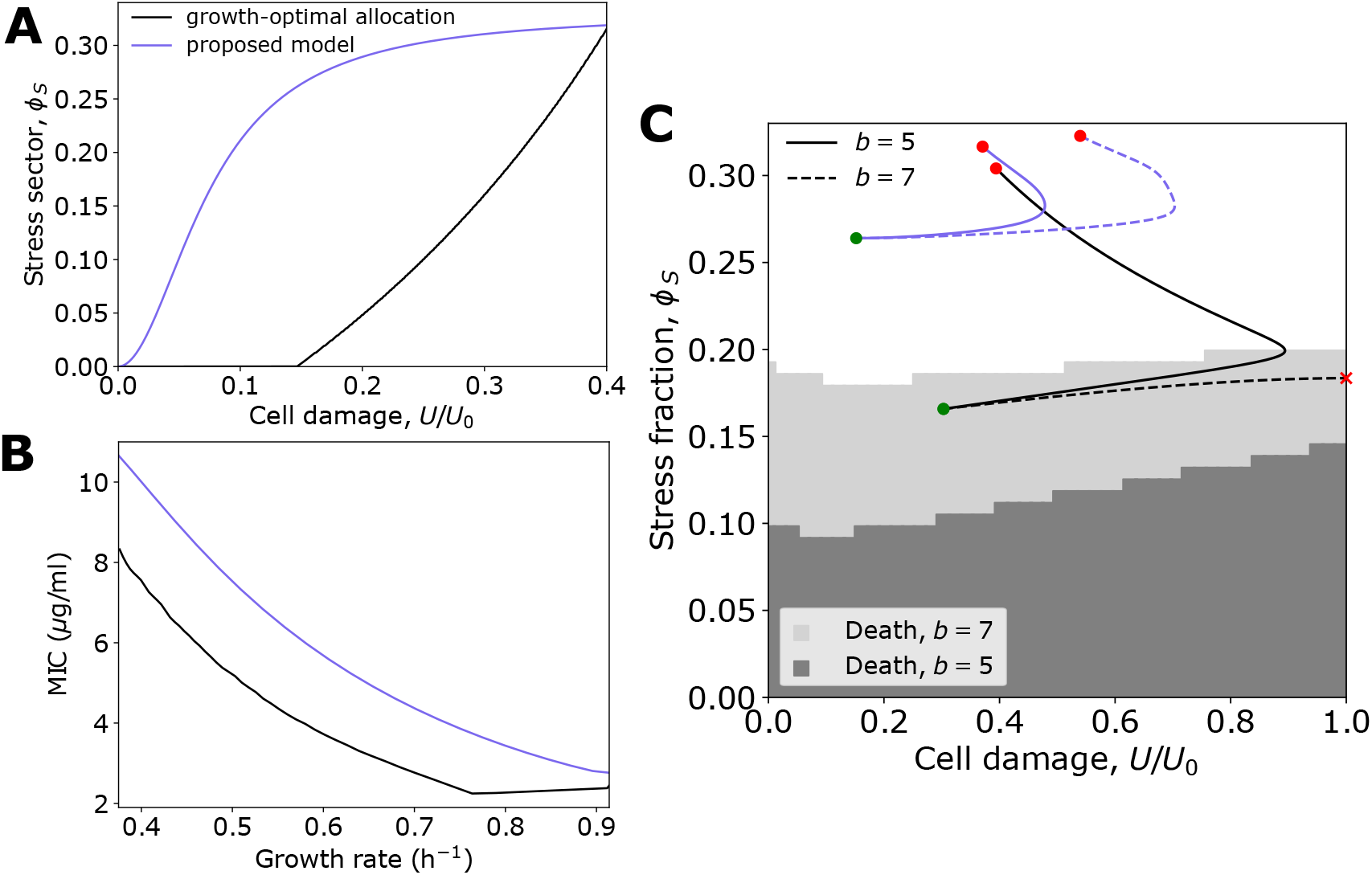
Non-growth optimal physiology increases bacterial survival. (A) Steady-state stress sector expression as a function of cellular damage for proposed model (blue) compared to growth-optimal allocation (black). For *U* ⦤ 0.15 growth-optimal allocation to stress protein expression remains at 0 because damage accumulation is prevented solely by growth dilution. (B) Predicted MIC for proposed model (blue) and growth-optimal allocation (black), with identical conditions as Figure 3A. (C) Cell outcome phase diagram for two values of *b*. Shaded regions indicate positions in phase space corresponding to cell death, whereas white indicates survival. Example trajectories from proposed model (blue) and growth optimal allocation (black), where dashed lines correspond to *b* = 7, and solid lines indicate *b* = 5. See Table 1 for a list of parameters.

To understand how the overexpression of *ϕ*_*S*_ results in an in-creased MIC, we can again consider cell state dynamics in the *ϕ*_*S*_-*U* phase space. With increasing antibiotic concentration *b*, the region in which the fixed point is accessible shrinks. Specifically, the boundary separating survival from cell death shifts upwards in the *ϕ*_*S*_-*U* plane (Figure 4C). Thus, by over-expressing *ϕ*_*S*_ at low levels of *b*, cell survival becomes more resilient to further increases in *b*. Conversely, in the growth-optimal model, stress sector expression remains significantly lower for the same *b* value, positioning it much lower in the phase diagram. Here, even slight increments in *b* lead trajectories to be absorbed towards the boundary *U* = *U*_0_ (see Figure 4C), causing cell death.

### E. Stochastic model of damage accumulation connects single-cell physiology to population-level behavior

Stress-induced cell death in an isogenic bacterial population is inherently stocahstic [2, 5]. When exposed to bactericidal antibiotics in particular, a significant fraction of bacteria die at sub-MIC concentrations, highlighting the stochastic nature of antibiotic killing. Furthermore, the fraction of surviving cells decreases in a dose-dependent manner [14, 38]. Our deterministic theory is limited to binary outcomes, namely a population either entirely survives or is completely eradicated, and so is unable to capture these phenomena. Therefore, to explain population-level behavior induced by bactericides, we must include the effects of stochasticity on cell death. In our model, cell death is ultimately determined by damage accumulation to the threshold level. Many factors can contribute to stochasticity in this process, including noise in gene expression, antibiotic import, and damage removal. As such, we choose to coarse-grain the noise in damage accumulation and reformulate the deterministic dynamics of *U* in terms of a Langevin equation, yielding

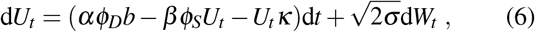

where *W*_*t*_ denotes a Wiener process with variance *σ*^2^. Example trajectories using this framework are given in Figure 5A, and show that cell death can occur even when the deterministic trajectory remains well below the critical damage threshold. Importantly, as the average damage level approaches the threshold, smaller deviations away from the mean are required for a cell to die, thus increasing the probability of death. More formally, (6) can be written in terms of a potential function, and the risk of death and survival fraction can be approximated as a function of antibiotic concentration in our model from the first-passage time of *U* above *U*_0_ (see Appendix D). Using the best-fit strain and antibiotic-specific parameters from the deterministic model, *σ* can be fit to yield good agreement between theory and data for experiments where survival fraction is not impacted by division rate (Figure 5B).

**FIG. 5.**
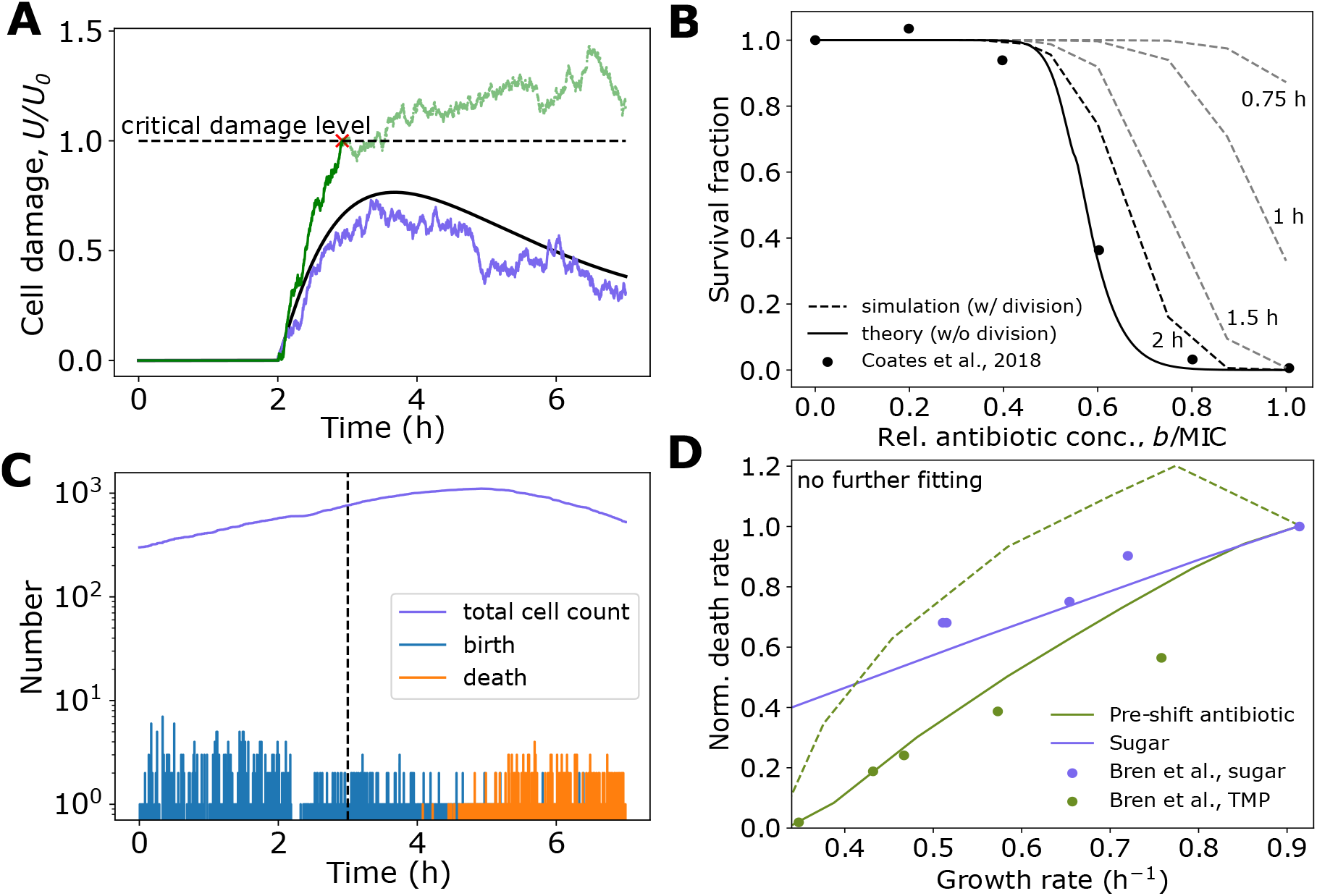
Stochastic model captures population-level behavior. (A) Representative trajectories of cell damage in response to antibiotic application at *t* = 2 h. At sub-MIC concentrations, the deterministic dynamics (black) remain below *U* = *U*_0_, but in the stochastic case, both survival outcomes are possible, with some cells surviving (blue) and others dying (green). (B) Survival fraction of a bacterial population exposed to various concentrations of antibiotic, relative to the MIC. Solid line denotes theoretical prediction from first-passage time of *U* above *U*_0_, dotted lines denote population simulation results for different durations of antibiotic exposure. Experimental data are from Ref. [38] and are of *E. coli* NCM3722 cells in LB exposed to ciprofloxacin, determined via plating efficiency. Model behavior is robust to parameter choice (see Fig. S7). (C) Population dynamics for bacteria exposed to antibiotics at *t* = 3 (dashed line), initialized with 300 cells. Following a division event, both resulting daughter cells were simulated. For each cell if *U ≥ U*_0_, the cell was removed, corresponding to cell death. (D) Death rate of *E. coli* in 10 *µ*g/ml nalidixic acid, defined as the inverse of the time required for 90% of the initial population (4000 cells) to be eliminated, as a function of pre-shift growth rate, normalized to the death rate in glucose. Solid lines indicate our proposed model, dashed line indicates growth-optimal resource allocation model. Data from Ref. [17]. See Table 1 for a list of parameters.

Although our stochastic model can capture the decrease in survival fraction with increasing antibiotic concentration seen experimentally, it does not take into account replication of surviving bacteria. Thus, to fully capture population-level behavior, we must both model single-cell death and division dynamics, which requires adding rules for division. To this end, we add a new sector, *X*, which regulates cell reproduction using rules for division and proteome allocation from our previous work (detailed in Ref. [40]). This allows us to simulate populations of cells in complex time-varying environments, with population dynamics governed by single-cell death and division events.

Using this stochastic multiscale model, we simulated single-cell trajectories for a population of cells, tracking the total number of living and dead cells over time in response to antibiotic application (Figure 5C). We found a significant portion of the population died at antibiotic concentrations below the MIC due to stochastic damage accumulation above the critical threshold, while a surviving subpopulation continued to grow and divide, in agreement with experimental observations [14]. Importantly, once *U* = *U*_0_, the cell dies, and damage cannot return to the mean value. Thus this absorbing boundary condition allows for the creation of two stable subpopulations defined by cell viability.

We calculated the fraction of surviving cells after simulating different time intervals of antibiotic exposure for increasing sub-MIC antibiotic concentrations, using the previous best-fit parameters. Not surprisingly, shorter exposure resulted in a higher survival fraction, with survival fraction stabilizing after several hours of exposure (Figure 5B). The dependence of exposure time on survival fraction is antibiotic-specific, because it is set by the dynamics of damage accumulation specific for each drug, thus highlighting the necessity of considering exposure time when assessing killing efficiency. The fraction of surviving cells decreased with increasing antibiotic concentration, however the survival fraction was always greater than that predicted by the single-cell theory (Figure 5B). This is because sub-MIC, surviving cells are able to continue to grow and divide, thus inflating the number of living cells.

### F. Population-level death rates are predicted by single-cell physiological state

To assess how the pre-shift environment and cellular stress response affects bacterial death dynamics, we used our multiscale model to simulate cell population dynamics in response to antibiotic exposure above the MIC. Here, to facilitate comparison with experimental data [17], we defined the death rate as the inverse of the time required for 90% of the initial population to be eliminated (1*/t*_90_). Interestingly, in all cases death rate decreased with decreasing growth rate, with cells pre-exposed to low levels of antibiotic having a lower death rate than those in a poor nutrient environment with the same growth rate (Figure 5D).

As with the predictions for the MIC, the reduction in death rate for cells pre-exposed to low levels of antibiotic is largely explained by the increase in stress protein expression causing a reduced rate of damage accumulation (Figure 3B). However, the decrease in death rate for cells in poor nutrient conditions is caused by a decrease in concentration of the antibiotic target, *ϕ*_*D*_. This can be understood by considering (1) and bearing in mind that, since we are considering gyrase-targeting antibiotics, *ϕ*_*D*_ ∝ *ϕ*_*R*_, and *ϕ*_*R*_ decreases with decreasing nutrient-imposed growth rate (Figure 2B). Consequently, slower-growing cells produce less damage (Figure 3B). As all cells initially have no damage, and thus no stress protein expression, and cell division is greatly reduced in all cases regardless of nutrient quality, differences in damage production dominate the dynamics of damage accumulation, and thus death rate. Importantly, we also simulated death dynamics using the growth-optimal resource allocation strategy, and found that death rates were significantly higher for this allocation strategy compared to our proposed model, and did not match the experimental data (Figure 5D). In fact, death rate increases at low pre-shift antibiotic levels in the growth-optimal model, as cells in this regime initially carry more damage, but do not induce a stress response (Figure 4A). This further supports the notion that bacteria utilize a non-growth optimal proteome allocation strategy in order to increase survival chances under antibiotic challenge.

### G. Differing timescales of stress exposure and proteome reallocation enable mutation-independent adaptation

With our model able to capture experimentally-observed growth rate, gene expression, and death dynamics, we then tested the model in more complex time-varying environments, focusing on pulsatile antibiotic exposure. We simulated a bacterial population growing in rich media subjected to repeated bactericide application. Interestingly, we found that for concentrations above the MIC of the initial population, population size recovered to its initial value after several pulses and then surpassed it as cells continued to proliferate (Figure 6A).

**FIG. 6.**
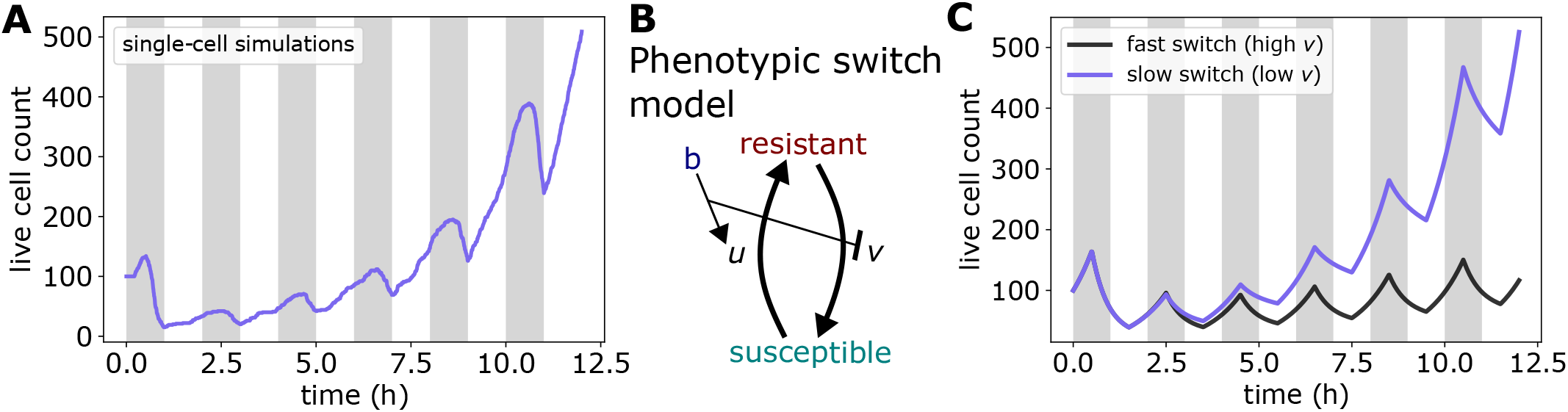
Proteome reallocation confers mutation-independent adaptation. (A) Population dynamics from single-cell simulations for bacteria under pulsatile antibiotic exposure above the MIC, initialized with 100 cells. Population recovers and surpasses initial size after 4 pulses. See Table 1 for a list of parameters. (B) Schematic depicting phenotypic switch model for population dynamics in time-varying environment. In response to antibiotic application (*b*), susceptible cells alter their gene expression to become more resistant at rate *u*, at the cost of a reduced growth rate. Upon antibiotic removal, cells switch back at rate *v*. See Appendix E for more details. (C) Phenotypic switch model can capture the adaptation to pulsatile exposure seen in (A).

These results identify a short-timescale, mutation-independent, adaptive response to bactericidal antibiotic exposure. Initially, antibiotic-induced damage accumulation occurs quickly, resulting in high rates of cell death. However, damage accumulation also causes bacteria to increase *ϕ*_*S*_ expression (Fig S4). As a result, cells which survive the initial pulse are better able to withstand subsequent exposure, resulting in an increased MIC and decreased death rate amidst future antibiotic challenge. Importantly, when antibiotics are removed, bacteria again reallocate their proteome to maximize growth, resulting in a decrease in *ϕ*_*S*_. Consequently, the observed adaptation is a result of the difference in timescales of antibiotic application and proteome reallocation. Specifically, when reallocation is slower than the time period of application, surviving cells will on average have a higher value of *ϕ*_*S*_ upon re-exposure compared to cells one period prior. As a result, the physiological state of surviving cells is better able to combat the next round of antibiotic application, thus yielding a higher survival fraction, in agreement with experimental observations [47].

Simulations in time-varying environments again highlight the importance of the physiological state of the cell in determining antibiotic killing efficacy. In such environments, this altering of physiological state can be modeled as a phenotypic switch between two subpopulations (Figure 6B), a framework which has been used successfully in pharmacodynamics (PD) to model resistance evolution [48], as well as in modeling bacterial persistence [6]. Critically, unlike previous work, here susceptibility is not altered via mutations or persister formation, but due to changes in gene expression (sector allocation) in growing cells (see Appendix E for full model description). Using this framework, we constructed a population-level description of our multiscale model which was able to reproduce the observed adaptive behavior (Figure 6C). Importantly, this adaptive behavior was mitigated when proteome reallocation after antibiotic removal was accelerated (Figure 6C), demonstrating that slow phenotypic switching can facilitate adaptation to pulsatile environments.

## III. DISCUSSION

We have developed a multiscale model for cell growth and death which connects extracellular antibiotic concentration and nutrient quality to bacterial physiology, allowing us to quantitatively capture the observed control of growth rate, death rate, minimum inhibitory concentration (MIC), and survival fraction across a wide range of environments. Although the model has been derived based on data from *E. coli* under replication-targeting bactericides, we expect the theoretical framework of proteome allocation theory and damage accumulation to generalize to other environmental stressors and other microorganisms. Using proteomics data in conjunction with MIC assays for a particular organism and drug pair, the various proteomic and kinetic parameters can be elucidated in our model, allowing for quantitative prediction of growth rate control and death rate in complex time-varying environments.

Our model reveals that cell death seldom occurs due to antibiotic levels exceeding the maximum physiological tolerance, but rather, cell survival hinges on the ability to transition to the appropriate physiological state. Consequently, the MIC of a bacterial population, typically assumed to remain constant unless raised by mutations [49], can be significantly altered by manipulating its physiological state through environmental changes. In addition, our model brings understanding to how changes in gene expression enable cellular adaptation in fluctuating environments. This is extremely pertinent over short timescales, when resistant mutations have yet to accumulate. As a result, our model has direct clinical relevance, as it allows for quantitative prediction and understanding of bacterial growth and death in time-varying nutrient and antibiotic environments at both the single-cell and population levels.

Furthermore, our model predicts that when exposed to bactericides, bacteria tend to overexpress stress response pathways at the expense of growth. This strategy enhances their resilience to future antibiotic challenge but comes at the cost of growth potential. This strategy highlights an important gene expression tradeoff that cells must make between growth and survival. Protection against environmental stress is expensive, as it requires synthesis of energy-consuming homeostatic mechanisms and repair processes [50]. Allocation towards such processes reduces the resources available for growth. Moreover, in fluctuating environments, rapid adaptation can confer a fitness advantage. As such, bacteria must continually respond to environmental changes to execute a program which balances the needs for both growth and survival.

Our modeling framework can easily be extended in future work to capture other environments and physiological contexts. The effects of dynamic stressors on drug-resistant bacteria, which constitute a serious global health problem [3, 4], can be studied by adding an additional proteome sector corresponding to the expression of resistance-conferring genes, or by altering the effective damage removal rate constant (*β*). In addition, our framework could be used to study the complex and nonintuitive effects of antibiotic combinations [1, 51, 52]. As our model is able to predict growth and death dynamics, it is particularly well suited to investigate temporal interactions between sequentially applied drugs in order to understand the effects of antibiotic-induced phenotypic changes on future drug applications.

## ACKNOWLEDGEMENTS

SB acknowledges support from the National Institutes of Health (NIH R35 GM143042), and the Shurl and Kay Curci Foundation. J.C.K. and S.B. designed and developed the study. J.C.K. carried out the simulations and analyzed the data. J.C.K. and S.B. wrote the article.

## APPENDIX A

### MODEL DERIVATION

To connect the physiological effects of bactericidal application to the population-level dynamics of cell death, we propose a coarse-grained process of damage accumulation and removal and connect it to other cell-level processes. Specifically, the dynamics of cell damage concentration *U* for a single cell can be expressed as 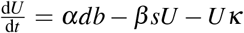 where *b* is the antibiotic concentration, which produces damage at a concentration specific rate *α* when bound to its target D, with concentration *d*. Damage is removed by stress proteins, with concentration *s*, at rate *β*, and is diluted by cell growth at rate *κ*. Cell death occurs when the damage level exceeds a critical concentration, *U* (*τ*_death_) = *U*_0_. In many conditions, cell density remains constant and total protein mass is proportional to the cell’s dry mass [53, 54], thus the concentration of each protein sector *i* can equivalently be expressed in terms of mass fraction, *ϕ*_*i*_. Therefore our equation for damage dynamics can be rewritten as

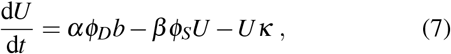

where the constants *α* and *β* are now mass fraction specific rates.

A significant number of known bactericides target the DNA gyrase, thus preventing replication. Steady-state proteomics data reveals that DNA gyrases are coregulated with ribosomal proteins under carbon, nitrogen, and translation limiting regimes [16]. Thus we assume such coregulation is maintained under DNA gyrase antibiotic challenge, such that its proteome fraction, *ϕ*_*D*_, is proportional to the R sector (*ϕ*_*D*_ = *νϕ*_*R*_). Therefore Eq. (7) becomes 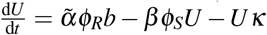 where 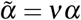

Following our previous model for dynamic proteome allocation [40], we can connect damage dynamics to changes in gene expression. Specifically, stress sector dynamics are given by

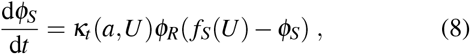

where *f*_*S*_(*U*) is the fraction of total cellular protein synthesis flux devoted to stress proteins and *κ*_*t*_ denotes the translational efficiency. We assume stress protein expression is solely dependent on the damage amount, such that 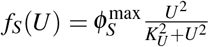 As before [40], *κ*_*t*_ is reduced by low levels of *a*. Importantly, *κ*_*t*_ is now also reduced by increases in *U* to capture the effects of damage on protein production. This reduction can be caused by direct inactivation of ribosomal proteins, as in the case of aminoglycosides [55], or be caused by indirect effects as is the case for quinolones, where DNA synthesis inhibition reduces protein production via a decrease in cellular DNA concentration [28]. Although we choose to model the effects of damage on growth through the translational efficiency, *κ*_*t*_, this can equivalently be thought of as a growth reduction caused by inactivation of translating ribosomes – indeed this yields a mathematically equivalent expression. Thus, 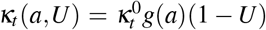 for 0 ≤ *U* ≤ 1,where the regulatory function *g*(*a*) is defined below. Similarly to Eq. (8), ribosomal sector dynamics are given by

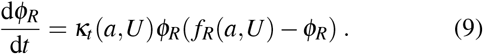

The rate of change of protein mass is proportional to *ϕ*_*R*_ [12], allowing us to define the growth rate of a single cell as *κ* = *κ*_*t*_(*a,U*)*ϕ*_*R*_.

The dynamics of the amino acid mass fraction, *a*, are given by the difference in the rate of nutrient import and conversion of nutrients to amino acids, and the rate of consumption via translation: 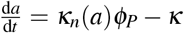 denotes the nutritional efficiency. Using the constraint 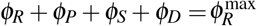, this becomes

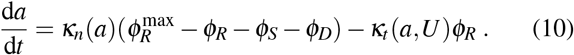

To make explicit the dependency of the efficiencies, *κ*_*n*_ and *κ*_*t*_, on *a*, we define two regulatory functions, *f* (*a*) and *g*(*a*), as given by Ref. [45]. Specifically, we assume 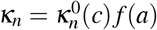 and 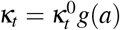 where 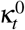 is a constant, and 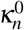 is a function of the extracellular nutrient concentration *c*. The regulatory functions are then:

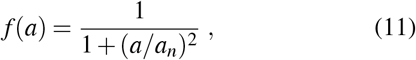

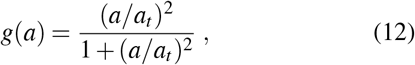

where translation becomes significantly attenuated for amino acid concentrations below *a*_*t*_, and the amino acid supply flux becomes significantly attenuated by feedback inhibition for *a* above *a*_*n*_.

Steady-state transcriptomic analysis reveals that ribosomal sector expression is reduced to allow for stress sector expression [17]. As such, the fraction of total synthesis capacity devoted to ribosomes, *f*_*R*_, is now a function of both *a* and *U*, specifically,

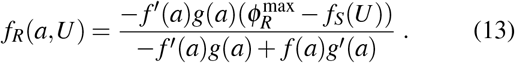

When *U* = 0, *f*_*R*_ is chosen to maximize translational flux at steady-state, thus maximizing growth rate [40].

## APPENDIX B

### COMPUTING THE MIC

As stated in the main text, the minimal inhibitory concentration (MIC) is defined in defined in terms of our model as:

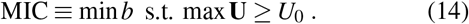

We take the critical threshold *U*_0_ to be constant, allowing Eq. (1) to be rewritten in terms of the normalized concentration, *Ũ*= *U/U*_0_, i.e.

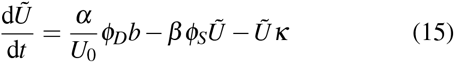

Thus, cell death occurs at *Ũ*= 1, and the fitting parameter *α/U*_*0*_, which represents the normalized damage production rate constant, determines survival.

As a result, we take *U*_*0*_ = 1 and used the normalized dynamics to solve the constrained optimization problem in Eq. (14). We use a simple line search method, in which an initial guess and jump distance are specified, the trajectory of U is solved via numerical integration of the coupled ODEs defined in Section 1, and then the guess and jump distance are updated each iteration based on the following criteria:

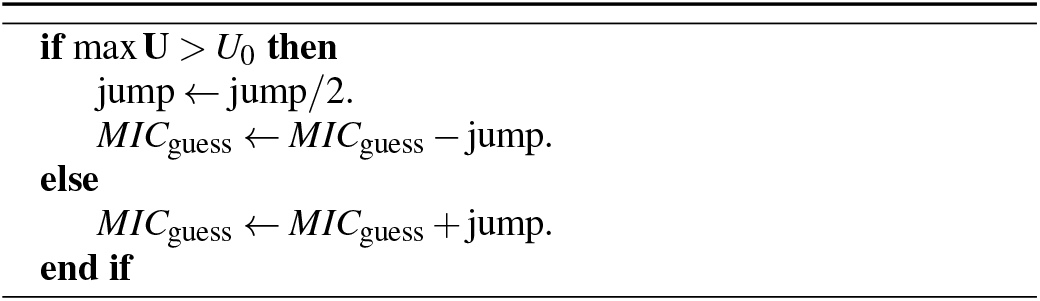

The algorithm typically converges within 10 iterations (Figure 7).

**FIG. 7.**
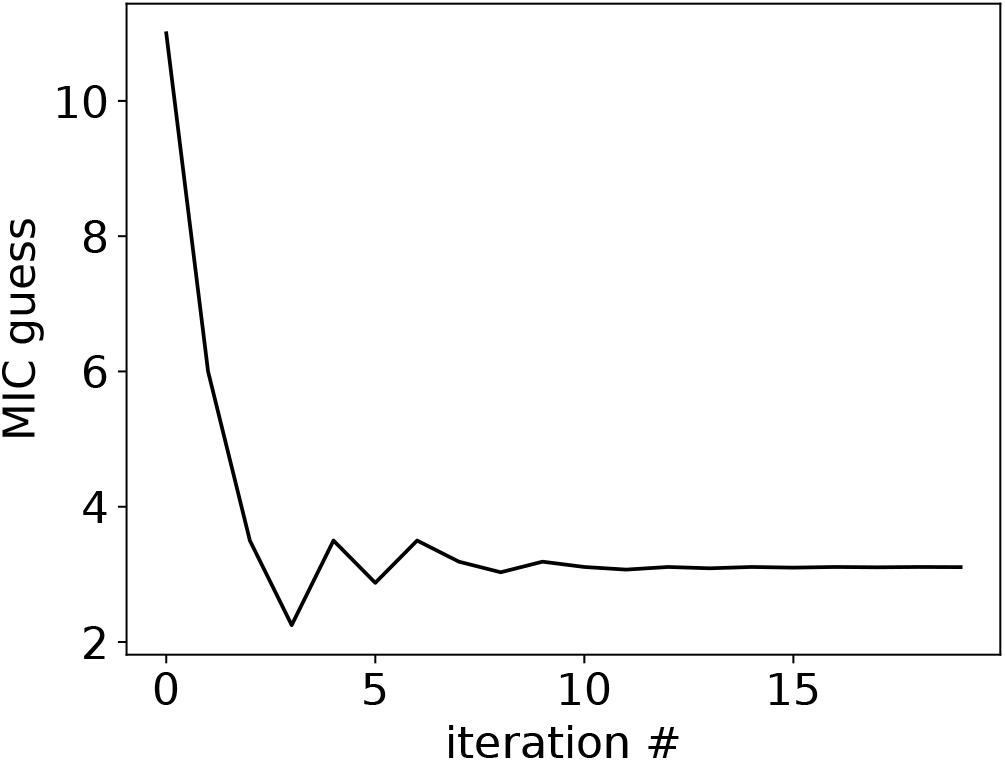
A simple line search method converges quickly to a solution for the MIC for a given environment. guess and jump distance are specified, the trajectory of **U** is solved via numerical integration of the coupled ODEs defined in Section 1, and then the guess and jump distance are updated each iteration based on the following criteria: The algorithm typically converges within 10 iterations (Figure 7).

## APPENDIX C

### NOTE ON BISTABILITY AND PROTEOME ALLOCATION

Antibiotic-induced growth bistability has been observed experimentally in antibiotic-resistant bacteria [4]. Interestingly, due to the nonlinear nature of target protein expression (*ϕ*_*D*_) as a function of damage (*U*), there is a predicted growth bistability in our model in very poor nutrient and high antibiotic environments (Figure 8, top). The stability of both solutions was confirmed by performing a linear stability analysis of the dynamical equations. In such conditions, faster growing cells carry less damage. Counterintuitively, this fast growth is achieved by reducing *ϕ*_*R*_ and *ϕ*_*D*_ expression, and is not mainly an effect of dilution (Figure 8, bottom). The increase in translational efficiency caused by the decrease in damage accumulation (caused by reduced target expression) outweighs the decrease in growth caused by reducing ribosomal expression. This bistable behavior is not predicted to occur in any of the environments in which we compare to experimental data.

## APPENDIX D

### DERIVATION OF RISK OF DEATH

Here we approximate the risk of death as a function of bactericidal antibiotic concentration using Kramer’s approximation for our resource allocation and damage accumulation model. The model dynamics for damage can be written as:

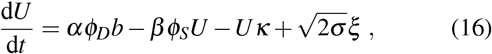

where *ξ* denotes Gaussian white noise. Following Ref. [39], this can be written in terms of a potential function *V* (*U, t*):

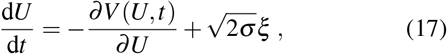

where the potential function is:

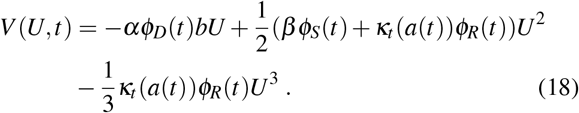

We model mortality as the first time when *U ≥* 1 after stepwise application of antibiotic with concentration *b*. Thus, death time is a first-passage time of *U*. To estimate the risk of death, i.e. hazard rate, we apply the Kramer approximation [56] for the mean first passage time:

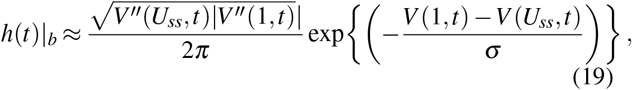

where *U*_*ss*_ denotes the quasi steady state of the system at time *t* after antibiotic application.

In the context of aging, it is usually of interest to calculate the hazard rate and survival function over time [57]. In contrast, here we are interested in understanding how the risk of death changes with increases antibiotic exposure. In this case, a cell is most likely to die when damage is at its maximum value, *U*^*∗*^, which our model predicts will occur shortly after antibiotic application. Thus to estimate the risk of death and survival fraction for a given antibiotic concentration, we define the hazard rate for a given value of *b* as:

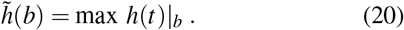

Thus using Eqs. (18), (19), and (20), we arrive at an expression for the hazard rate in terms of our model parameters:

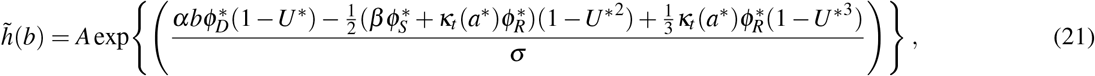

where the prefactor *A* is given by:

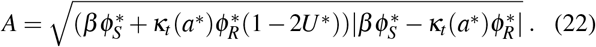

Importantly *a*^*∗*^, 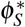, and 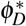 are the amino acid mass fraction and proteome allocation fractions of the S and D sectors when *U* = *U*^*∗*^, and thus depend on *b*. Eq. (21) can be solved numerically and is plotted in Figure 9.

**FIG. 8.**
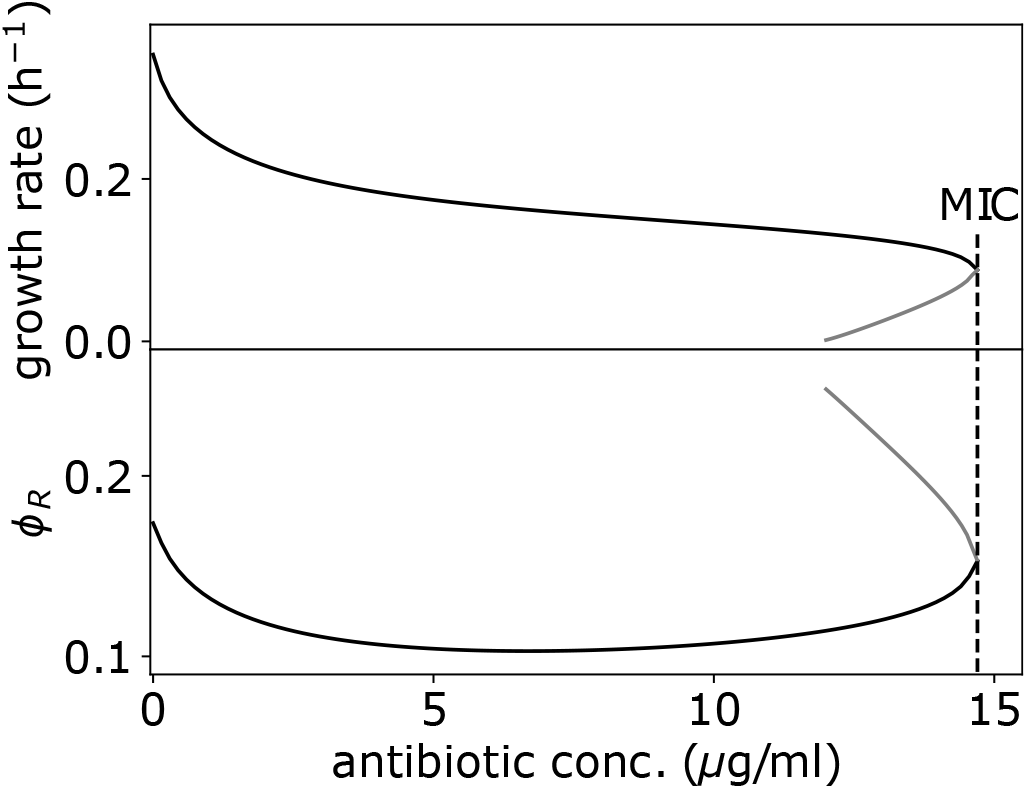
Predicted growth rate (top) and ribosomal allocation (bottom) bistability. Increased ribosomal expression (gray) corresponds with decreased growth rate (gray).

**FIG. 9.**
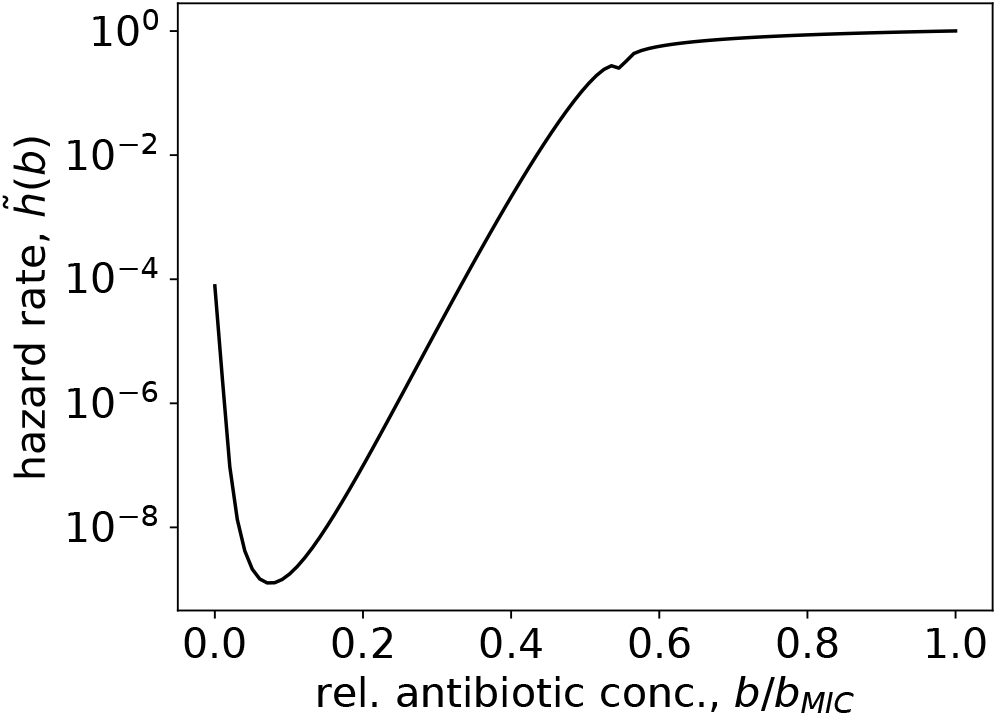
Maximum hazard rate as a function of applied antibiotic concentration.

This hazard function is related to the survival function, which is the cumulative probability of remaining alive, through the relation:

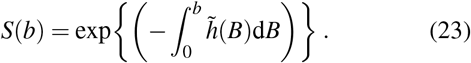

Eq. (23) can be integrated numerically to predict survival fraction as a function of antibiotic concentration, and can be fit well to experimental data (Figure 5).

## APPENDIX E

### PHENOTYPIC SWITCHING MODEL OF POPULATION GROWTH AND DEATH IN PULSATILE ENVIRONMENTS

Here we write a continuum description of population-level growth and death dynamics in response to pulsatile antibiotic exposure. Cells grown in rich media are initially highly susceptible to bactericide application, characterized by a low MIC and high death rate [17]. In response to damage accumulation caused by antibiotics, bacteria alter their gene expression profile, yielding an increase in *ϕ*_*S*_. As a result, those that survive the initial pulse are better able to withstand subsequent exposure, resulting in an increased MIC and decreased death rate, at the expense of a reduced growth rate. This altering of physiological state can be modeled as a phenotypic switch between two subpopulations, a framework which has been used successfully in pharmacodynamics (PD) to model resistance evolution [48], as well as in modeling bacterial persistence [6]. Critically, unlike previous work, here susceptibility is not altered via mutations or persister formation, but due to changes in gene expression (sector allocation) in growing cells.

PD curves model the effect of drug concentration on net growth rate of bacteria [48]. In the absence of antibiotics, bacteria grow at rate *κ*_max_, which is set by the nutrient environment. When antibiotics levels are well above the MIC, bacteria are killed at rate *κ*_min_ (*κ*_min_ *<* 0). The growth curve for a given nutrient environment can then be defined as [48]:

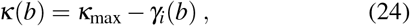

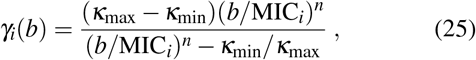

where here *i* = [*S, R*] denotes susceptible cells (*P*_*S*_) or cells with increased MIC due to an altered physiological state (*P*_*R*_), with MIC_*R*_ *>* MIC_*S*_, and where *n* is the Hill coefficient. We use the notation *P*_*i*_ to stress the fact that they denote two different physiological states of isogenic replicating bacteria. The drug-dependent growth rate can then be used to predict overall population-level dynamics, *P*(*t*), from the two subpopulations:

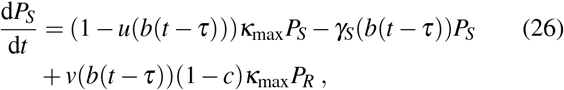

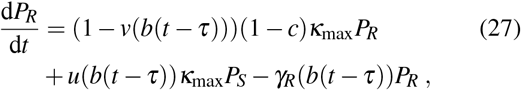

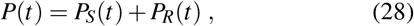

where here *κ*_max_ is the susceptible bacterial subpopulation maximum net growth rate, and *c* is the growth cost of altering gene expression to increase resistance. *u*(*b*) and *v*(*b*) denote the rate of phenotype switching from *P*_*S*_ to *P*_*R*_, and *P*_*R*_ to *P*_*S*_, respectively, corresponding to proteome reallocation. For constant antibiotic exposure, this model reduces to a form which is mathematically equivalent to the Type II persister model presented by Balaban et al. [6], however here we model two subpopulations of growing cells and do not consider the formation of persisters. In Eqs. (26) and (27), both *u*(*b*(*t − τ*)) and *v*(*b*(*t − τ*)) are functions of the antibiotic concentration with delay *τ*, such that cells switch from *P*_*S*_ to *P*_*R*_ at rate *u*_max_ with some delay in response to antibiotic-induced damage, and switch back from *P*_*R*_ to *P*_*S*_ at rate *v*_max_ with some delay when antibiotics are removed to maximize growth. Specifically,

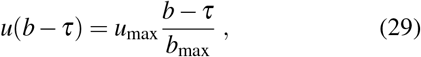

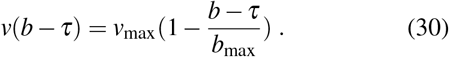

When antibiotics are pulsed at concentrations above the MIC in a static population which cannot adapt, the population will perish (Figure 10A). If surviving cells can alter their physiological state to become more resistant to antibiotics, after several pulses cells are able to adapt and proliferate (Figure 10B). Critically, this adaptation occurs only when the timescale of returning to the susceptible phenotype (set by *v*_max_) is less than the time period of antibiotic pulsing. This population-level description closely matches what is seen in our single-cell simulations (Figure 6).

**FIG. 10.**
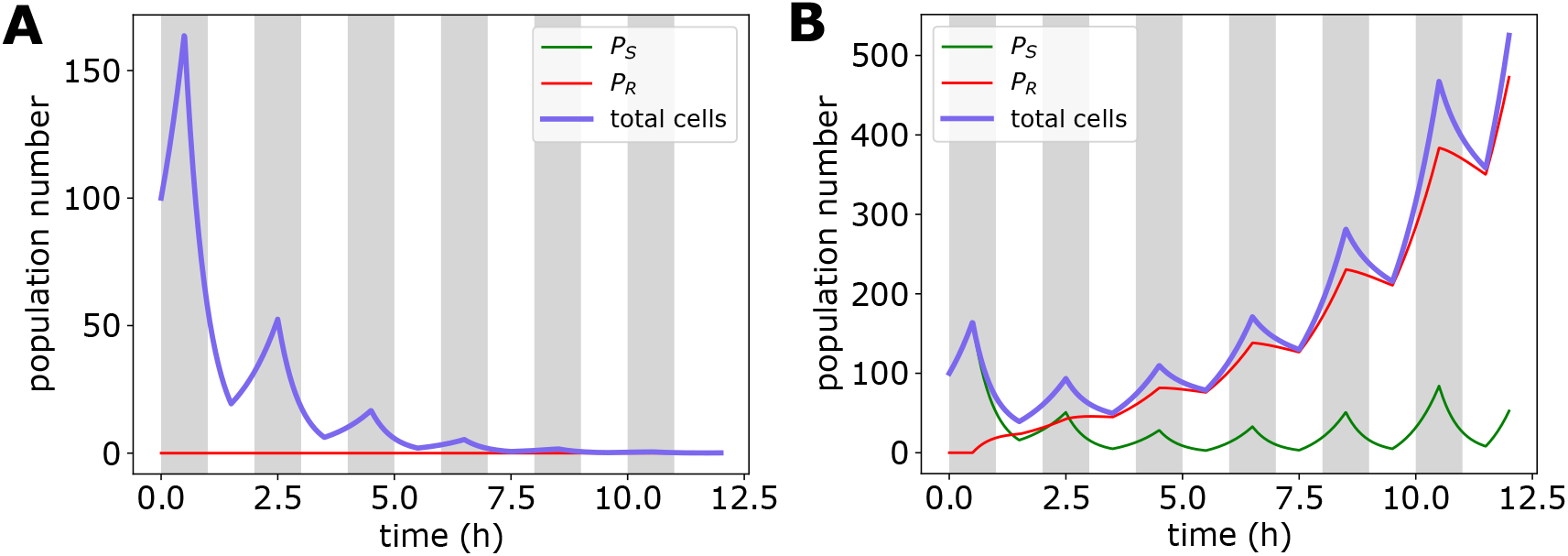
Population model reproduces adaptation to pulsatile antibiotic exposure seen in our single-cell resource allocation and damage accumulation model. (A) Static population does not survive antibiotic exposure above the MIC. (B) Adaptive population can survive and proliferate by altering its physiological state.

