## Supplemental Material for "Gene expression tradeoffs determine bacterial survival and adaptation to antibiotic stress"

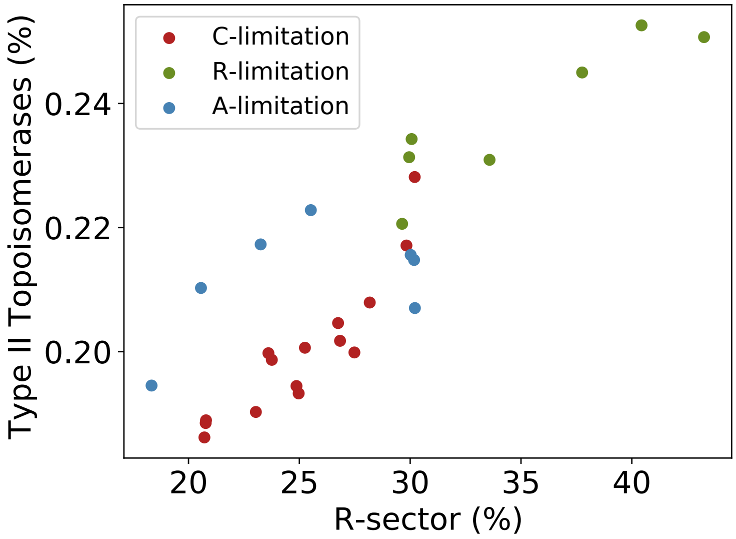


Fig. S1: **Coregulation of quinolone targets with R-sector.** Both naladixic acid and ciprofloxacin are classified as quinolone antibiotics [1], which target the type II topoisomerases DNA gyrase (composed of gyrA and gyrB) and topoisomerase IV (composed of parE and parC). Mori et al. [2] measured the proteome composition (defined in terms of mass fraction) of *E. coli* under carbon, nitrogen, and translational limitation. The abundance of the type II topoisomerases are shown as a function of the ribosomal (R) sector. For the three regimes tested, they exhibit a strong positive correlation (correlation coefficient 0.87), indicating coregulation of topoisomerases with the ribosomal sector.

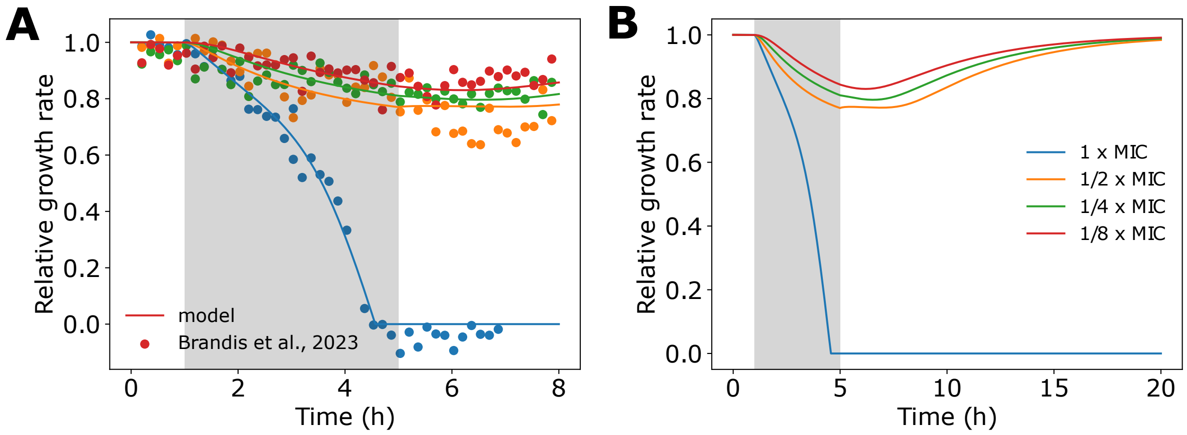


Fig. S2: **Growth rate dynamics during antibiotic pulse.** (A) Model parameters obtained by fitting growth rate data at 1 x MIC (blue dots). Orange, green, and red lines correspond to model predictions (no further fitting), and agree well with experimental data (dots). (B) When antibiotics are removed, growth rate recovers to the preshift rate after several hours if concentration is below the MIC.

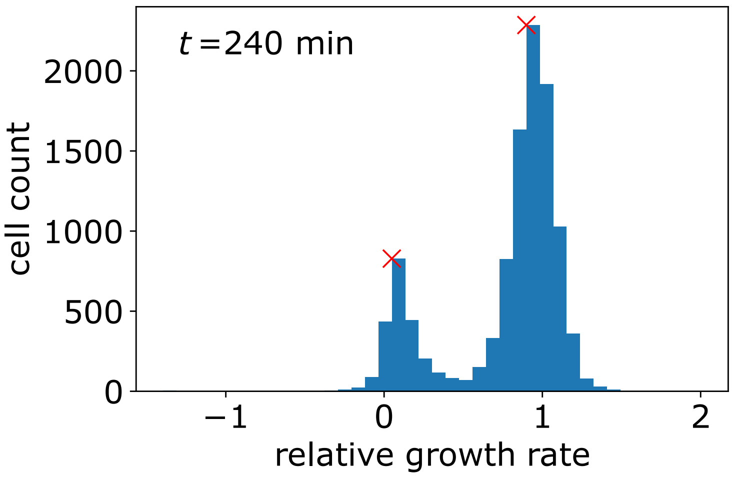


Fig. S3: **Bactericidal application produces two sub-populations of cells.** Example relative growth rate distribution of *E. coli* cells after 4 hours of 0.016 mg/L Ciprofloxacin exposure, which produces two sub-populations of cells; growing and dead. Data from Ref. [3]. The growth rate corresponding to the maximum of each distribution (red x) was identified in 10 min intervals using the signal.find_peaks function from scipy, yielding the growth rate dynamics of the surviving population plotted in Figure 1 in the main text.

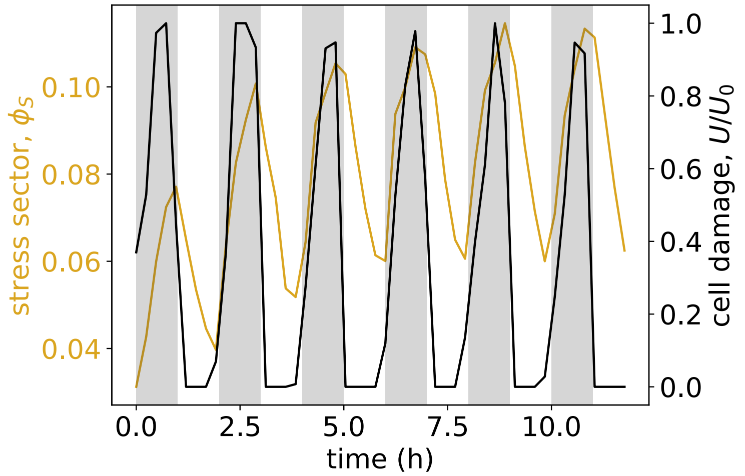


Fig. S4: **Bacteria alter physiological state in pulsatile environments to adapt.** Population-averaged dynamics of cell damage and stress sector expression from single-cell simulations for bacteria under pulsatile antibiotic exposure above the MIC, corresponding to the population dynamics shown in Figure 6A in the main text. See Table 1 for a list of parameters.

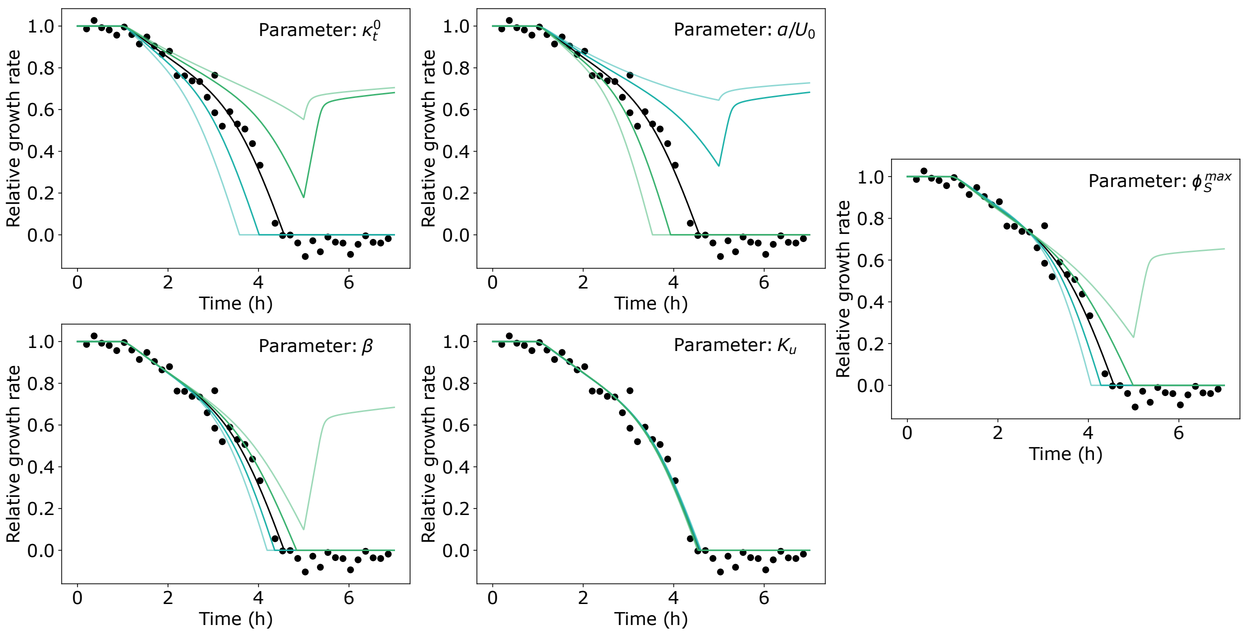


Fig. S5: **Assessing effects of parameter choice on growth control behavior.** Each of the five strain or bactericide-specific fitting parameters corresponding to the setup detailed in Figure 1E were varied by + (green) or − (blue) 20% of their best-fit value to assess the effects of parameter choice on model behavior. Darker colored lines denote +/−10%. An antibiotic pulse was applied from $t = 1$h to $t = 5$h, thus trajectories which increase after 5 hrs correspond to survival. See Table 1 for list of best-fit parameter values. Data from Ref. [3].

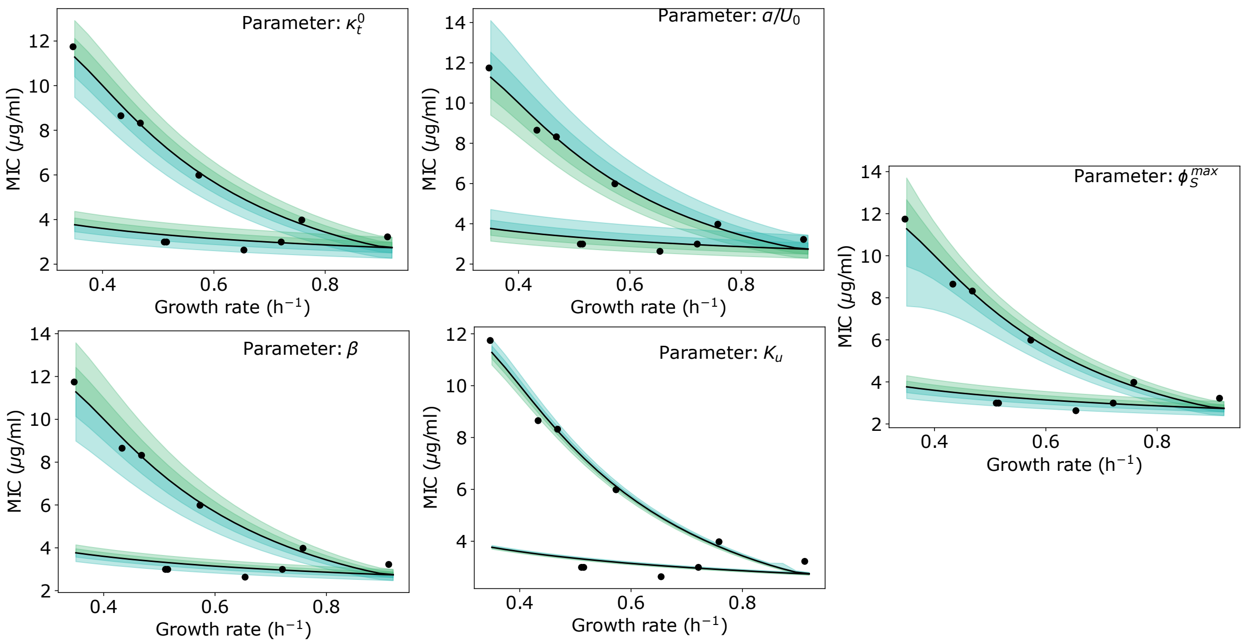


Fig. S6: **Predicted MIC trends are robust to parameter choice.** Each of the five strain or bactericide-specific fitting parameters corresponding to the setup detailed in Figure 3A were varied by + (green) or − (blue) 20% of their best-fit value to assess the effects of parameter choice on model behavior. Darker shading denotes +/−10%. Qualitative model behavior remains unchanged regardless of parameter choice. See Table 1 for list of best-fit parameter values. Data from Ref. [4].

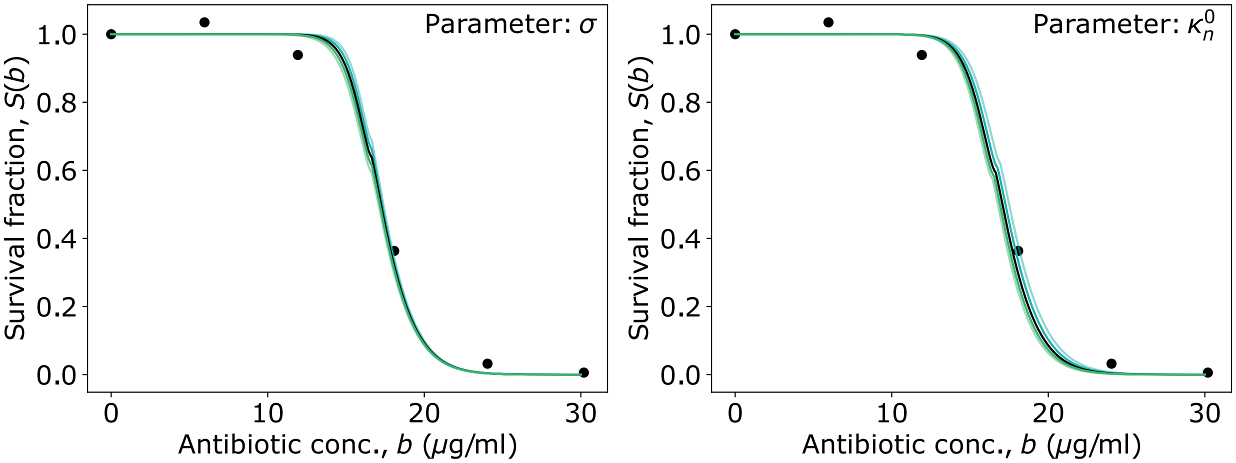


Fig. S7: **Assessing effects of parameter choice on survival fraction.** By fitting to experimental data as shown in Figures S5 and S6, the five strain or bactericide-specific fitting parameters were determined. Keeping these fixed, we varied the two environment-specific fitting parameters by + (green) or − (blue) 20% of their best-fit value to assess the effects of parameter choice on model behavior. Darker shading denotes $+/-10\%$. Predicted model behavior changes little despite large deviations in parameter choice. See Table 1 for list of best-fit parameter values. Data from Ref. [5].
